# Worldwide survey reveals lower susceptibility of African *Aedes aegypti* mosquitoes to diverse strains of Zika virus

**DOI:** 10.1101/342741

**Authors:** Fabien Aubry, Daria Martynow, Artem Baidaliuk, Sarah H. Merkling, Laura B. Dickson, Claudia M. Romero-Vivas, Anubis Vega-Rúa, Isabelle Dusfour, Davy Jiolle, Christophe Paupy, Martin N. Mayanja, Julius J. Lutwama, Alain Kohl, Veasna Duong, Alongkot Ponlawat, Van-Mai Cao-Lormeau, Richard G. Jarman, Cheikh T. Diagne, Oumar Faye, Ousmane Faye, Amadou A. Sall, Louis Lambrechts

## Abstract

Zika virus (ZIKV) is a flavivirus mainly transmitted to humans through the bite of infected *Aedes aegypti* mosquitoes. First isolated in Uganda in 1947, ZIKV was shown to circulate in enzootic sylvatic cycles in Africa and Asia for at least half a century before the first reported human epidemic occurred in 2007 on the Pacific island of Yap, Micronesia. Subsequently, larger ZIKV outbreaks were recorded in French Polynesia and other South Pacific islands during 2013-2014. In 2015, ZIKV reached Brazil from where it rapidly spread across the Americas and the Caribbean, causing hundreds of thousands of human cases. The factors that have fueled the explosiveness and magnitude of ZIKV emergence in the Pacific and the Americas are poorly understood. Reciprocally, the lack of major human epidemics of ZIKV in regions with seemingly favorable conditions, such as Africa or Asia, remains largely unexplained. To evaluate the potential contribution of vector population diversity to ZIKV epidemiological patterns, we established dose-response curves for eight field-derived *Ae. aegypti* populations representing the global range of the species, following experimental exposure to six low-passage ZIKV strains spanning the current viral genetic diversity. Our results reveal that African *Ae. aegypti* are significantly less susceptible than non-African *Ae. aegypti* across all ZIKV strains tested. We suggest that low susceptibility of vector populations may have contributed to prevent large-scale human transmission of ZIKV in Africa.

## Main text

Zika virus (ZIKV) is a mosquito-borne virus (genus: *Flavivirus*; family: *Flaviviridae*) mainly transmitted among humans through the bite of infected *Aedes aegypti* mosquitoes [1]. ZIKV was first isolated in 1947 from the serum of a sentinel rhesus monkey in the Zika Forest of Uganda [2]. Subsequently, serological evidence and virus isolation from humans and mosquitoes revealed ZIKV circulation in enzootic sylvatic cycles both in Africa (African ZIKV lineage) and Asia (Asian ZIKV lineage). For at least half a century following its discovery, ZIKV did not cause any recorded human epidemic, as less than 20 human cases were documented between 1947 and 2006 [1,3].

The first ZIKV human outbreak occurred in 2007 on the Pacific island of Yap in the Federated States of Micronesia, Oceania where ZIKV (Asian lineage) infected 73% of the islanders [4]. ZIKV outbreaks were then reported in French Polynesia [5] and other South Pacific islands during 2013-2014 [6]. Whereas human ZIKV infection is usually asymptomatic or results in a self-limiting mild illness, for the first time it was associated with severe neurological complications such as congenital microcephaly and Guillain-Barré syndrome [7,8]. In 2015, ZIKV reached Brazil and rapidly spread across South, Central, and North America and the Caribbean [3], causing an estimated 583,451 suspected and 223,477 confirmed cases between 2015 and 2017 [9].

To date, the factors that promoted the rapid emergence, dispersal and apparent increase in pathogenicity of ZIKV in the Pacific and the Americas are not fully understood. They may include a number of mechanisms, including viral mutations increasing transmission from humans to mosquitoes, enhancing fetal microcephaly, and modulating the host immune response [10-14]. Reciprocally, the lack of major epidemics in regions such as Africa and Asia, where conditions are seemingly favorable, is still largely unexplained. It has been hypothesized that this may reflect higher levels of background immunity conferred by cross-protective antibodies against ZIKV-related viruses [15].

Besides ZIKV evolution or contrasted human background immunity, the potential contribution of vector population genetic diversity to ZIKV epidemiological patterns may have been overlooked [15]. Earlier studies using field-derived *Ae. aegypti* populations demonstrated significant variation in ZIKV vector competence at different geographical scales [16-21]. However, none of these studies alone considered the entire geographical range of *Ae. aegypti*. Likewise, the ZIKV strains previously tested did not span the extent of ZIKV genetic diversity. Here, we established dose-response curves for eight field-derived *Ae. aegypti* populations from across the global distribution of the species (Table 1) exposed to six low-passage ZIKV strains (Table 2) representing the current breadth of viral genetic diversity (Figure 1).

**Table 1.** *Aedes aegypti* colonies included in this study. The country of origin, locality, year of collection and number of generations spent in the laboratory prior to this study are indicated.

| Country | Locality | Year | Generations |
| --- | --- | --- | --- |
| Thailand | Kamphaeng Phet | 2013 | 13-14 |
| Cambodia | Phnom Penh | 2015 | 8-9 |
| Colombia | Barranquilla | 2017 | 2-3 |
| French Guiana | Cayenne | 2015 | 5-6 |
| Guadeloupe | Saint François | 2015 | 6-7 |
| Gabon | Lopé | 2014 | 9-11 |
| Uganda | Zika | 2016 | 4-5 |
| Cameroon | Bénoué | 2014 | 7-8 |

**Table 2.** ZIKV strains included in this study. The country of origin, source of viral isolation, year of collection and passage history prior to the study are indicated. TS: *Toxorhynchites splendens* mosquitoes; C6/36: *Aedes albopictus* cells; Vero: green monkey kidney cells; SM: suckling mouse brains.

| Country | Source | Year | Passage history |
| --- | --- | --- | --- |
| Thailand | Human serum | 2014 | TS-1; C6/36-2; Vero-1 |
| Cambodia | Human serum | 2010 | Vero 2; SM-1 |
| Philippines | Human serum | 2012 | TS-1; C6/36-2; Vero-1 |
| French Polynesia | Human serum | 2013 | C6/36-3 |
| Puerto Rico | Human serum | 2015 | Vero-2; C6/36-2 |
| Senegal | Pool of *Aedes* spp. and *Mansonia* spp. | 2015 | C6/36-1 |

**Figure 1.**
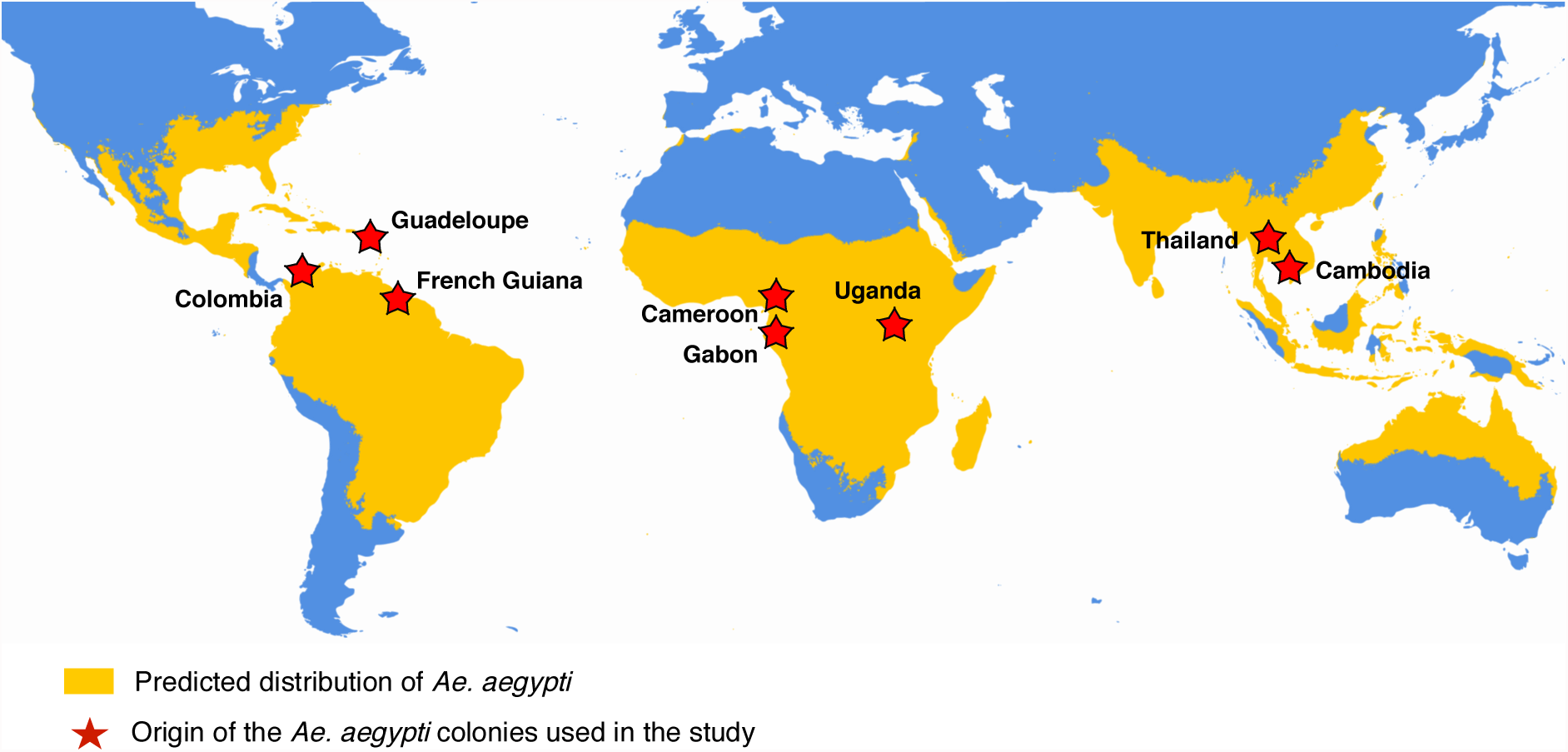
Geographical origin of field-derived *Ae. aegypti* colonies used in this study. The locations of origin of the colonies are indicated by the red stars, overlaid with the approximate global distribution of *Ae. aegypti* adapted from Kraemer et al. [22].

**Figure 2.**
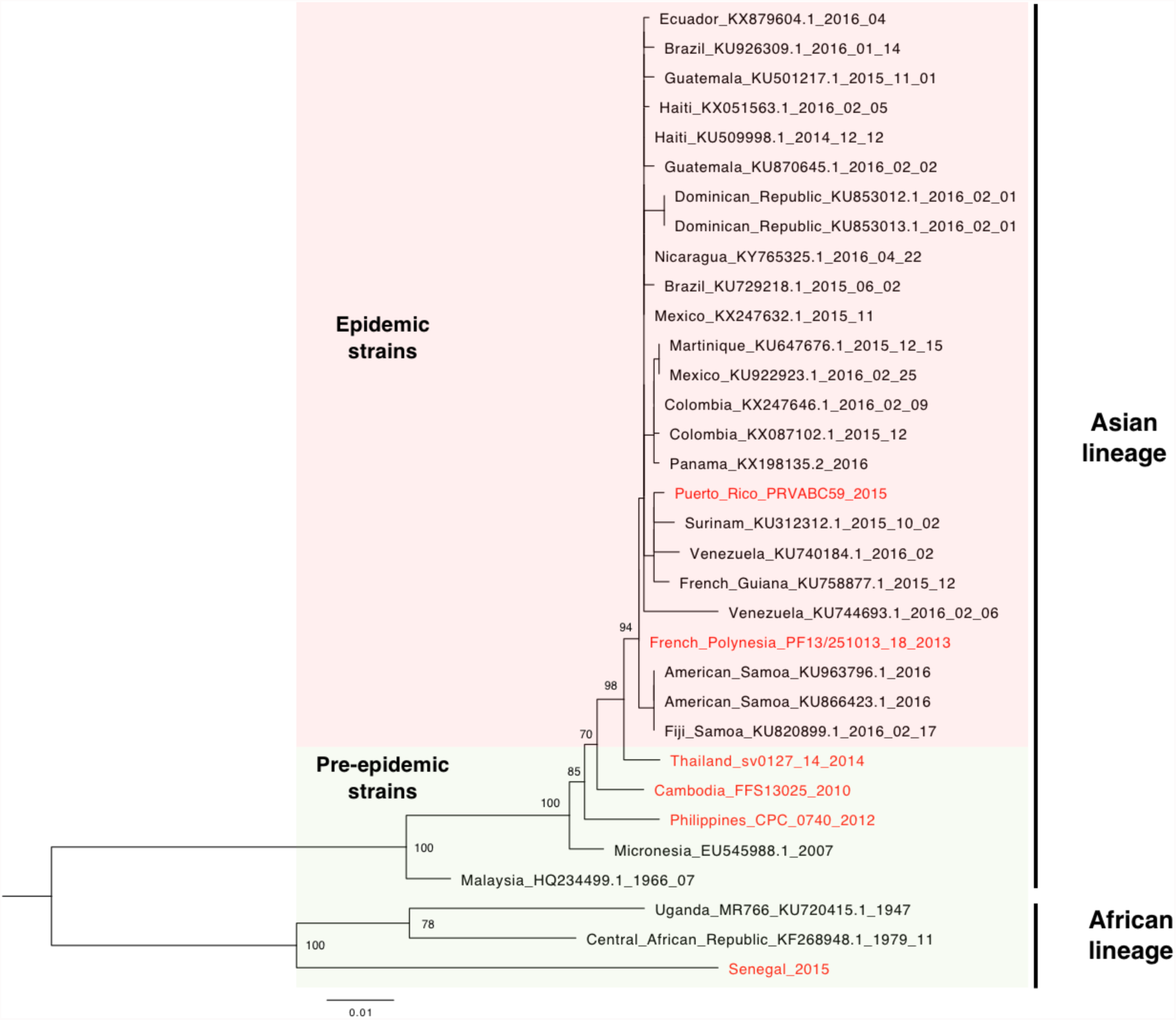
Phylogenetic position of ZIKV strains used in this study. Maximum-likelihood tree based on ZIKV envelope gene sequences shows ZIKV strains used in this study (in red font) among a background set of ZIKV strains (in black font) spanning the current viral genetic diversity. The light red background represents ‘epidemic’ ZIKV strains isolated during a human outbreak. The light green background represents ‘pre-epidemic’ ZIKV strains isolated in non-epidemic context, with the exception of the Micronesia strain, which caused a human outbreak in 2007. Bootstrap support values are shown at the relevant nodes and the scale bar represents the number of substitutions/sites.

Field-derived *Ae. aegypti* colonies from Africa (Cameroon, Uganda, Gabon), America (Colombia, Guadeloupe, French Guiana) and Asia (Thailand, Cambodia) were simultaneously challenged by oral exposure to one of six ZIKV strains, including one strain of the African lineage (Senegal 2015), three pre-epidemic (Philippines 2012, Cambodia 2010, Thailand 2014) and two epidemic (French Polynesia 2013, Puerto Rico 2015) strains of the Asian lineage. The artificial infectious blood meal consisted of a mixture of washed erythrocytes and viral stock (three different concentrations) produced in C6/36 cells, supplemented with ATP. Upon exposure, blood engorged mosquitoes were maintained at 28°±1°C under 12h dark:12h light cycle and 70% relative humidity. After 7 days of incubation, whole mosquito bodies were processed for RNA extraction and RT-PCR to estimate ZIKV infection prevalence.

A total of 3,113 female *Ae. aegypti* were individually scored for ZIKV infection and the overall prevalence was 52.7%. Multivariate logistic regression showed that the infection status depended on a three-way interaction between infectious dose, virus strain and mosquito population (*p* = 0.0238), indicating that the dose-response curves differed significantly among virus-population pairs (Figure 3).

**Figure 3.**
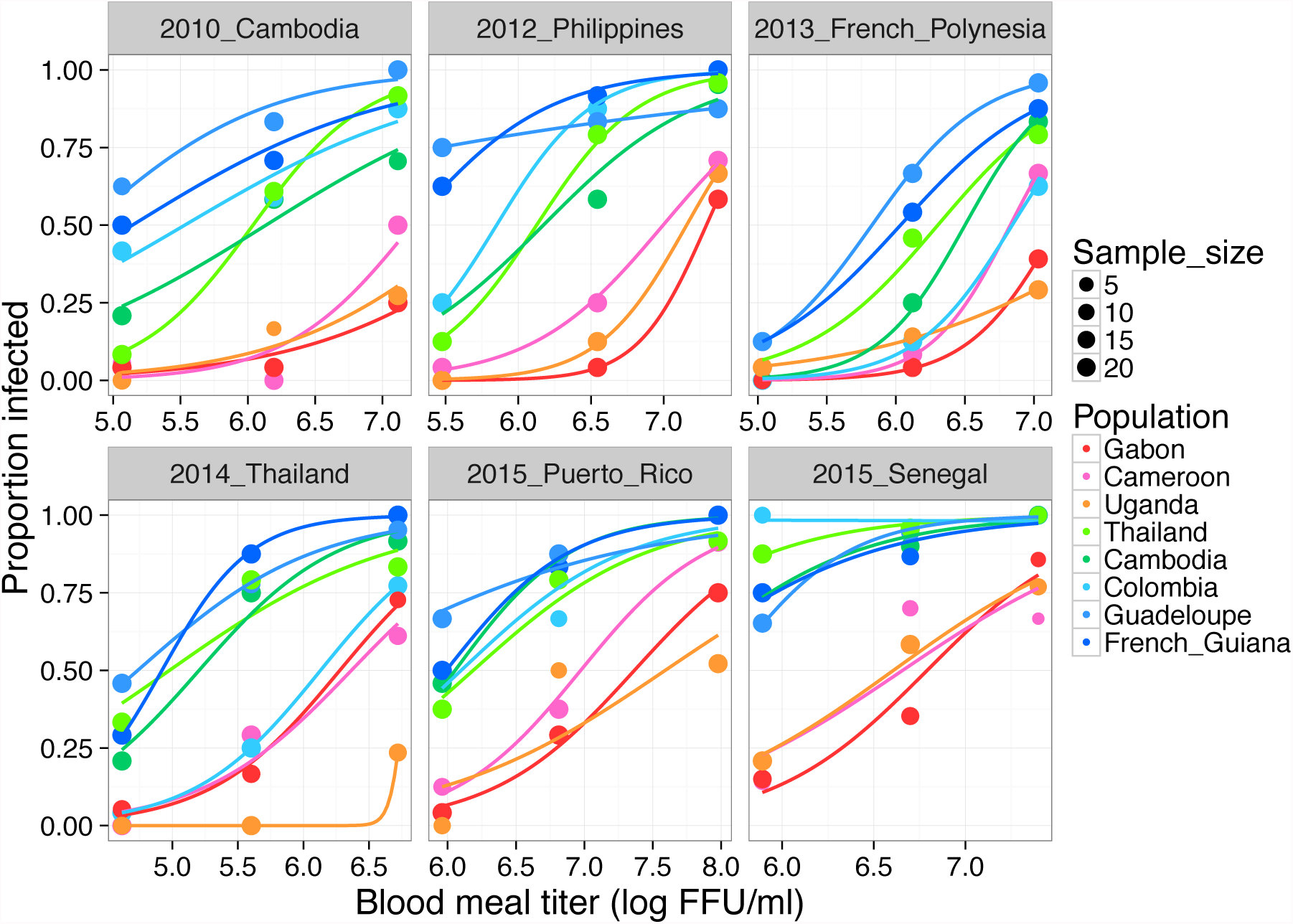
Dose-response curves of eight field-derived *Ae. aegypti* colonies challenged by six low-passage ZIKV strains. The percentage of ZIKV-infected mosquitoes 7 days post oral challenge is shown as a function of the blood meal titer. Each panel represents a different ZIKV strain. Lines are logistic regressions of the data, color-coded by mosquito population. FFU: focus-forming units.

When mosquito populations were nested within their continent of origin (Asia, Africa, Americas) in the statistical model, there was a strong effect of the continent (*p* < 0.0001), which was mainly driven by the significantly lower susceptibility of the three African mosquito populations. This was reflected in their 50% oral infectious dose (OID_50_), defined as the blood meal titer resulting in 50% prevalence. Across ZIKV strains, the OID_50_ estimates obtained from the dose-responses curves ranged from 6.3 to 8.1 log_10_ FFU/ml for the three African populations whereas they ranged from 4.7 to 6.8 log_10_ FFU/ml for the five non-African populations (Figure 4).

**Figure 4.**
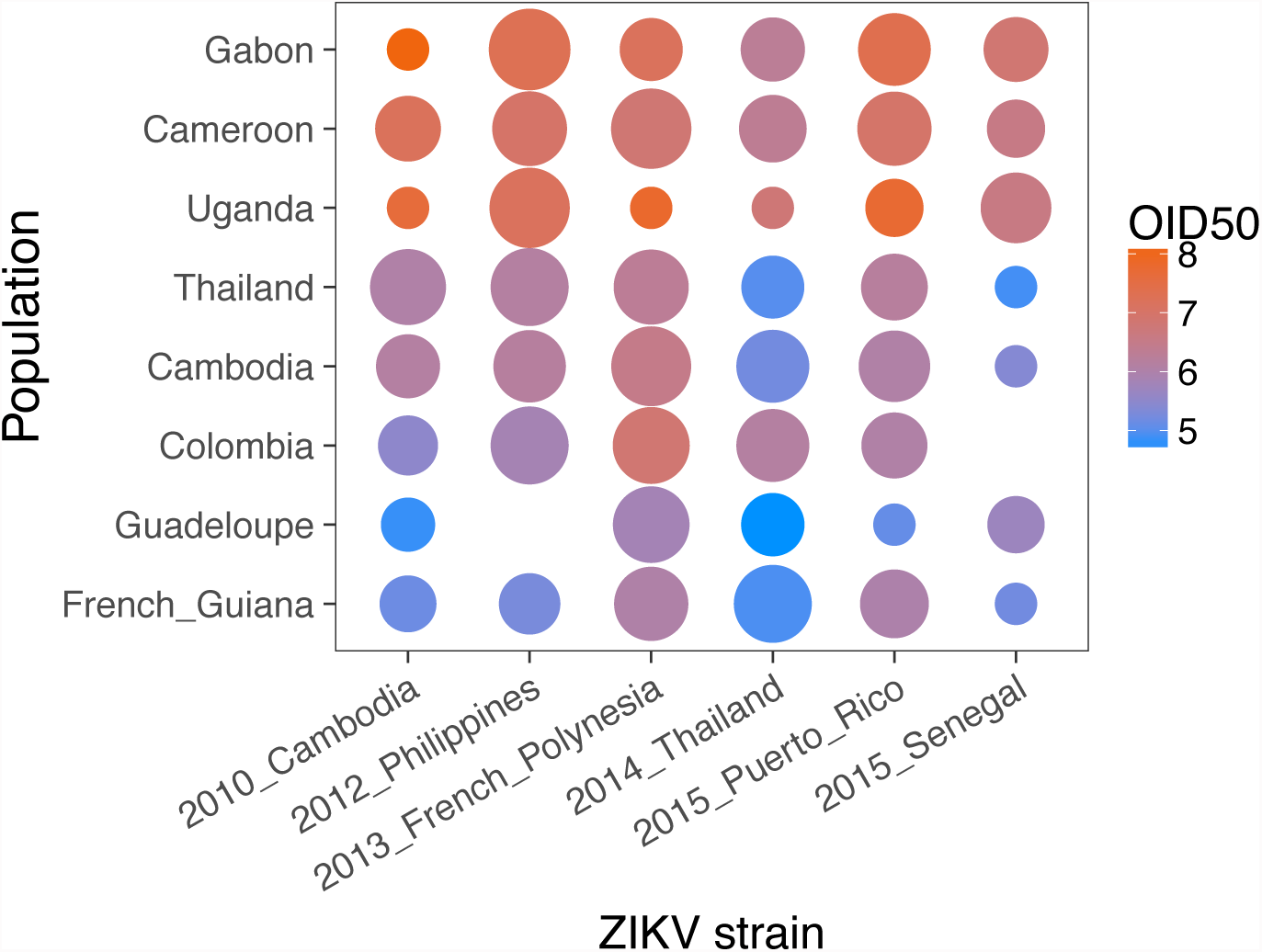
African *Ae. aegypti* are less susceptible to ZIKV infection. OID50 estimates are shown for each virus-population pair on a color scale in log_10_ FFU/ml. The size of the dot is inversely proportional to the size of the confidence interval of the OID_50_ estimate. When the size of the confidence interval could not be estimated it was arbitrarily set to 2 log_10_ units. Lack of a dot means that the OID_50_ could not be estimated with the data.

Our data indicate that *Ae. aegypti* colonies of African origin are, overall, less susceptible to all ZIKV strains tested than *Ae. aegypti* colonies of non-African origin. Oral susceptibility does not directly translate into vector competence because the latter also depends on subsequent steps of virus dissemination and transmission. However, it is tempting to speculate that the lower susceptibility of African *Ae. aegypti* populations may have contributed to prevent large-scale human ZIKV outbreaks in Africa. Exceptions are Gabon where ZIKV presumably circulated in the human population in 2007 but was likely transmitted by *Ae. albopictus* [23] and Cabo Verde where a human ZIKV outbreak occurred in 2015 off the coast of West Africa [24]. Conversely, our data fail to provide an explanation to the lack a major human epidemic in Asia because the two Asian *Ae. aegypti* populations tested had similar levels of ZIKV susceptibility as American populations. The clear dichotomy observed in ZIKV susceptibility between our African and non-African *Ae. aegypti* colonies mirrors the two main genetic clusters of global *Ae. aegypti* populations [25]. Elucidating the genetic basis of this natural difference in ZIKV susceptibility could help to unravel the mechanisms of ZIKV acquisition by mosquitoes.

## Acknowledgements

This work was primarily funded by the European Union’s Horizon 2020 research and innovation programme under ZikaPLAN grant agreement no. 734584. It was partially supported by the European Union’s Horizon 2020 research and innovation programme under ZIKAlliance grant agreement no. 734548.

## Author contributions

Designed and coordinated the study: FA, LL. Performed the experiments: FA, DM, AB, SHM, LBD. Contributed biological materials: CMRV, AVR, ID, DJ, CP, MNM, JJL, AK, VD, AP, VMCL, RGJ, CTD, OumF, OusF, AAS. Analyzed the data: FA, LL. Wrote the paper: FA, LL.

